# Clonal reproduction as a double-edged sword: antagonistic and synergistic effects on the warmed population maintenance of alpine species across successional stages of plateau zokor disturbance

**DOI:** 10.64898/2026.07.18.739372

**Authors:** Hai-Tao Miao, Shuai Jiang, Zhenhua Zhang, Shou-Li Li

## Abstract

A central question in biodiversity conservation under global change is whether species maintain viable populations under both mammal disturbance and climate warming. This requires demographic studies that integrating vital rates’ responses to mammal disturbance and climate warming across an entire life to. Using Integral Projection Models parameterized with demographic data, we found the population growth rates of *Thermopsis lanceolata* under both ambient and warming conditions, initially decreased on new mounds, further declined on seminew mounds, but eventually exceeded initial levels on old mounds. This stage-dependent responses were largely driven by clonal reproduction (i.e., clonal production and/or ramet size distribution), which emerged as both the most sensitive vital rate and the primary contributor to variation in population growth rates across recovery stages. Additionally, we found that the combined effects of plateau zokor disturbance and warming on population growth rates of new mounds was greater than the sum of their individual effects, leading to population decline on new mounds. Such synergistical effects was mainly due to a larger decrease of ramet size distribution. These findings suggest that multifactorial experiments are important for biodiversity research, rather than merely adding single effects on population dynamics. In addition, clonal reproduction may be a key vital rate for population maintenance under global change.

## INTRODUCTION

Disturbance from small fossorial mammals poses a growing threat to the persistence of plant populations worldwide (Huntly and reichman 1994, Reichman and Seabloom 2002, Niu et al. 2020, Asefa et al. 2024). Concurrently, climate warming may further exacerbate the impacts of such disturbances on population viability (Jiang et al. 2013, Panetta et al. 2018, Miao et al. 2025). Assessing plant population viability under disturbance and warming requires demographic studies that integrate species’ vital rates (e.g. survival, growth, and reproduction) across the life cycle in response to both disturbance and warming (Easterling et al. 2000, Caswell 2001). However, most studies have focused on the effects of a single factor, either disturbance or warming, on individual performance (Andersen and Macmahon 1985, Hansen et al. 2023, Zhang et al. 2024) and community dynamics (Reichman and Seabloom 2002, Hu et al. 2017, Wang et al. 2020), while largely overlooking their combined effects on population demography (Wolfe-Bellin and Moloney 2000). This knowledge gap limits our ability to accurately predict population dynamics under disturbance in both current and warming climates, and to develop effective conservation and management strategies in the context of ongoing global change.

Whether a plant population can persist after disturbance largely depends on its ability to resist and recover from such events (i.e., demographic resilience; Capdevila et al. 2020). Recent demographic studies have developed a theoretical framework to quantify demographic resilience using transient population dynamics (Stott et al. 2011), and demonstrated that demographic resilience can be captured by three key components: resistance, compensation, and recovery time (Figure 1; Capdevila et al. 2022). However, most applications of this framework have focused on natural populations, with limited empirical evidence from populations experiencing disturbance (Stott et al. 2010). Consequently, the scarcity of demographic data collected during post-disturbance recovery has hampered our understanding of the mechanisms underlying demographic resilience. Moreover, climate warming may further modify post-disturbance recovery processes (Miao et al. 2025), yet its effects on demographic resilience remain poorly understood. Therefore, assessing population viability during post-disturbance recovery under current and future climate warming is crucial for understanding how warming shapes the recovery trajectories of disturbed populations.

**Figure 1.**
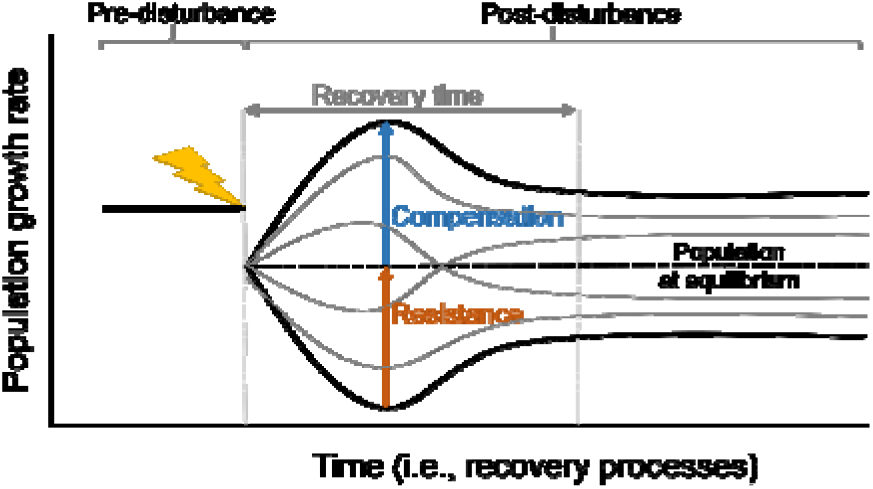
Demographic resilience of a natural population following a punctual disturbance (lightning bolt) can be decomposed into three key components: compensation, resistance, and recovery time. At a stable equilibrium population is expected to change in size at a constant rate (black dashed line). The growth rate of a population after a disturbance may vary (depicted by grey lines) depending on how the population structure (i.e. the proportion of individuals at different size within the population) is affected by the disturbance. Resistance, the ability to avoid a decline in population growth rate following a disturbance, is quantified as the inverse of a population’s decrease following a disturbance relative to its undisturbed states (i.e. stable population structure). Low resistance indicates a large decline in population growth rate relative to a stable population. Compensation, the ability to increase relative to the population growth rate following a disturbance, is measured as the population’s increase with respect to an undisturbed population. Large increases in population growth rate indicate high compensation, whilst small increases indicate low compensation. Recovery time, the period required for the population to return to a stable structure after a disturbance.

Another challenge hindering our ability to predict outcomes is that the individual effects of disturbance and warming on demography (e.g. certain vital rate or population viability) often differ from their combined effect (Cote et al. 2016, Zettlemoyer 2023). When disturbance and warming act on demography separately, their combined effect may be dominated by one factor (i.e. equal to the effect of one of dominant factor, “dominance”), or determined by a threshold calculated from the impacts of disturbance and warming (i.e. equal to the sum of their individual effect, “additive”) (Figure 2). Alternatively, disturbance and warming can interact either synergistically or antagonistically. If individual responses are in same direction, synergy occurs when the combined impact of disturbance and warming exceeds the threshold, while antagonism occurs when their combined effect falls below the threshold (Figure 2a, b). However, when the individual responses are in opposite directions, the sum of the individual effects is used as the threshold to differentiate between synergy and antagonism (Figure 2c). Moreover, synergistic responses of vital rates to disturbance under a warming climate may amplify their impacts on population performance, while antagonistic responses may weaken their impacts. Thus, mechanistically understanding these interactions is crucial for accurately predicting the effects of disturbance and warming on population dynamics.

**Figure 2.**
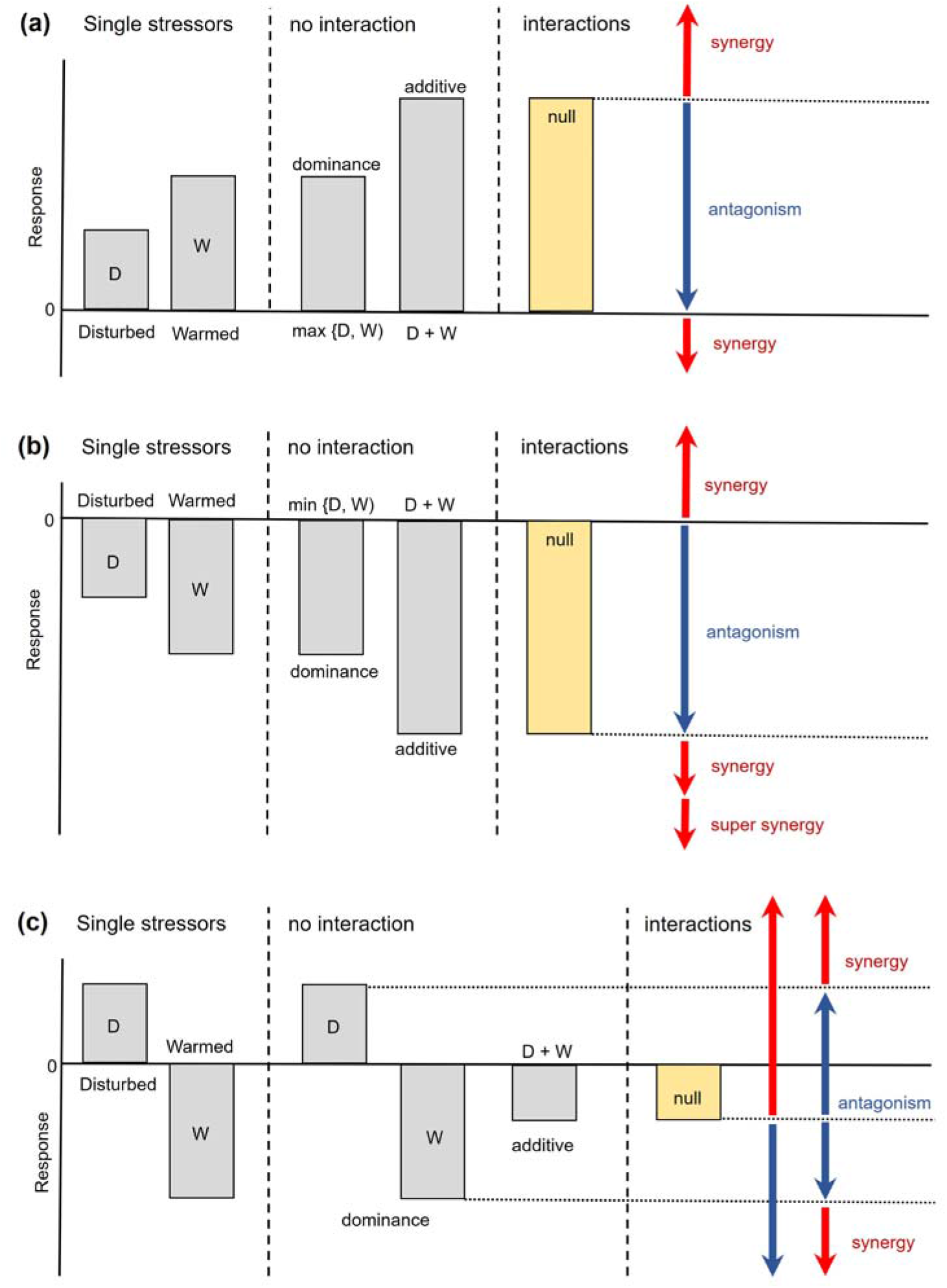
Defining demographic antagonism and synergy between multiple stressors (i.e. disturbance and warming). (a, b) Two stressors (D and W) impact a demographic response in the same direction when acting separately. Their combined effect could simply be equal to the effect of one of the two stressors, i.e. a dominance effect, or be additive, i.e. the sum of the two stressor effects. Alternatively, stressors can interact either antagonistically or synergistically. An additive expectation provides the threshold for distinguishing between these interactions. ‘super-synergy’ occur that push populations below their minimum viable population size does extinction risk become nonnegligible (Brook et al. 2008). (c) Two stressors have opposing effects on a demographic response. Here, some authors use a null model to define interaction types, as done in this study (Folt et al. 1999, Crain et al. 2008), while others advocate using the effects of each stressors to classify interactions (Piggott et al. 2015).

Alpine grasslands are considered among the most vulnerable ecosystems to disturbance and climate warming (Seddon et al. 2016, Nguyen and Liou 2019). Mound-building activity by plateau zokor (*Myospalax baileyi*) is the one of the common disturbances on alpine grasslands of the Tibetan Plateau, where the annual temperature has risen at twice the global average rate over the past 50 years (Yao et al. 2022). Therefore, exploring the consequences of plateau zokor disturbance under climate warming for population persistence of the Tibetan Plateau grasslands is curtail for informing the management of such critical susceptible alpine ecosystems. Moreover, such findings can provide early warning signals for the conservation of other ecosystems. Previous studies on the Tibetan Plateau grasslands have found that disturbance from plateau zokors and warming shifted plant community composition (Liu et al. 2018, Niu et al. 2020). However, it remains unclear how the persistence of alpine plant populations on the Tibetan Plateau will be affected by plateau zokor disturbance and climate warming.

Here, we examine how plateau zokor disturbance and climate warming affects the population dynamics of a dominant alpine herb, *Thermopsis lanceolata* R. Br., on the Tibetan Plateau grassland. To quantify the demographic responses both warming and disturbance, we parameterized integral projection models (IPMs) using demographic data collected in 2022 and 2023 (Caswell 1989, Easterling et al. 2000). We used these models to test the following hypothesis: (1) plateau zokor disturbance and climate warming negatively affect individual vital rates, thereby reducing population growth rate; (2) The combined effect of plateau zokor disturbance and climate warming on population growth rate will be greater than the sum of their individual effects, due to synergistic effects on vital rates; (3) Thus, climate warming will weaken resistance and prolong recovery time of a population following plateau zokor disturbance. If these hypotheses are supported, our results will highlight that climate warming may intensify the demographic consequences of plateau zokor disturbance, with important implications for predicting the persistence of alpine plant populations on the Tibetan Plateau.

## MATERIALS AND METHODS

### Study area and experimental design

The study was conducted at the at the Qinghai Haibei National Field Research Station of Alpine Grassland Ecosystem, located in the northeastern of the Tibetan Plateau, Qinghai Province, China (Figure 3a). The climate is characterized by a long, cold winters and a short, cool summers. The mean annual temperature is −1.1℃, with the highest monthly temperature occurring in July (10.4℃) and the lowest monthly temperature in January (−14.6℃). The mean annual precipitation is 488 mm, with over 80% falling as rain during the growing season from May to September (Wang et al. 2020). The site is typical alpine meadow dominated by perennial herbaceous plants, including *Thermopsis lanceolata*, *Carex alatauensis*, and *Potentilla anserine* (Liu et al. 2018). The plateau zokor (*Eospalax baileyi*), a typical subterranean rodent endemic to the Tibetan Plateau (Thomas 1911, Andrew et al. 2010), acts as a keystone ecosystem engineer in this alpine grassland (Niu et al. 2020). By continuously excavating burrows and pushing soil to the surface, they form mounds that actively reshape the microtopography of the alpine grasslands (Figure 3b). When plateau zokor populations become overly excessively dense, there cumulative disturbances can exceed the ecosystem’s recovery capacity, resulting in biodiversity losses and even grassland degradation. The study site was fenced annually in summer (from June to September) to avoid big animal disturbance but was grazed annually in winter by yaks and sheep (from October to May) (Fuller et al. 2017).

**Figure 3.**
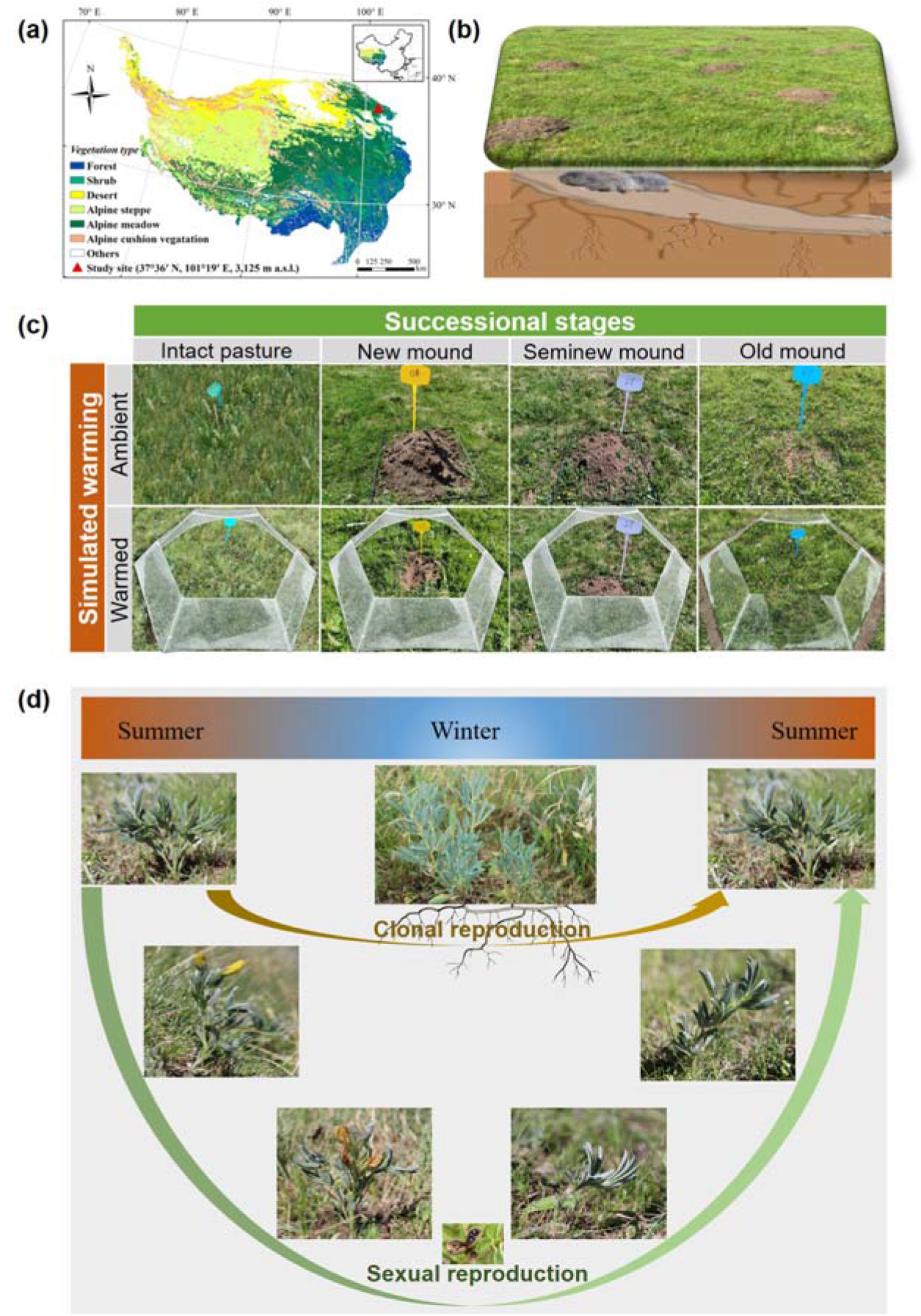
Demography of the alpine herb *Thermopsis lanceolata* under two experimental factors (plateau zokor disturbance and simulated warming) in an alpine meadow on the Tibetan Plateau. (a) Geographic location of the study site on the Tibetan Plateau. (b) Mound landscape formed by the burrowing activity of plateau zokor (*Eospalax baileyi*) in the alpine meadow. (c) Experimental design for different succession stages (intact pasture, new mound, seminew mound, and old mound) of plateau zokor mound under ambient and warmed conditions. (d) Life cycle of the study species reproducing both clonally and sexually.

To assess the effects of plateau zokor disturbance and climate warming on the population dynamics of alpine herbaceous plant, a two-factorial completely randomized design was established in July 2022 within an area of 8 ha (Figure 3c). Plateau zokor disturbance was classified into four successional stages—undisturbed pasture, new mound, seminew mound, and old mound—based on mound age, soil compaction, and vegetation characteristics (Table S1; Su et al. 2015, Tang et al. 2021). The warming treatment include ambient and warmed conditions. To simulate climate warming across succession stages, transparent hexagonal open-top chambers (OTCs) were installed. Each OTC had an internal diameter of 2.2 m and a height of approximately 0.65 m. This design is commonly used in the remote alpine ecosystems to passively elevated temperature (Elmendorf et al. 2012, Prager et al. 2022). The OTCs increased the mean air temperature by ∼2℃ and soil temperature in the top 5 cm layer by ∼1℃ (Prager et al., 2022), consistent with the predicted warming of 2℃ by 2050 (IPCC 2023). All the treatments were randomly assigned within study area, with three 1 m × 1 m plots per treatment (Figure S1a, b). To prevent grazing disturbance during winter, all plots were enclosed with mesh cages, effectively excluding yaks and sheep (Figure S1c).

### Study species and demographic census

To assess the effects of plateau zokor disturbance and climate warming on the population performance of alpine plant, we selected a perennial herb *Thermopsis lanceolata* (Figure 3d). The plant is a legume with grey-green leaves and erect tapering panicles of pale yellow, pea-like flowers. It sprouts new leaves annually, regrows in late April, flower in July, set fruit in August, and its aboveground parts wither in winter. Recruitment occurs both via seed germination and clonal propagation through horizontally spreading underground rhizomes that produce new ramets. The plant is widely distributed across Asia and Europe (Royal Botanic Gardens 2026) and serves as a forage plant in its early growth stages, as well as a medicinal species in traditional remedies (Flora of China Editorial Committee 1994).

To examine the effects of plateau zokor disturbance and climate warming on demographic processes, annual censuses were conducted in August 2022 and 2023 for alpine herb *T. lanceolata*. In the first census, we measured individual plant height and stem diameter, and counted the number of branches. We also recorded the reproductive status of each individual and counted the number of flowers produced by those that were reproductive. Upon the first measurement, we tagged each individual with a stainless-steel label, and recorded its Cartesian coordinates within the plots to allow tracking through time. In 2023, we checked the survival of all tagged individuals and remeasured plant size and aforementioned reproductive characteristics on surviving ones. At that time, we also located and measured the new recruits within our plots, including both new seedlings and newly produced ramets. Because distinguishing seedlings from new ramets is not feasible without destructive sampling, new recruits were identified as seedlings if they had a stem diameter below 0.5 cm, a height below 10 cm, and lacked branches and flowers; all other recruits were considered new ramets. Since adult plants usually produce numerous ramets and tracing the ‘parent’ of new ramets is not possible without destructive sampling, we assumed that clonal reproduction is size-dependent. Specifically, the new ramets produced by an individual was assumed to be linearly related to its size (Verburg et al. 1996). Accordingly, the number of new ramets produced per individual was estimated by multiplying its size (*x*) by the ratio *n*⁄∑ *x*, where *n* is the total number of new ramets in the current year and ∑ *x* is the sum of the sizes of all individuals in the previous year within each plot (Li et al. 2013, Li et al. 2015). In total, we tracked 6,778 individuals across all plots.

### Demographic rate estimates

To evaluate the effects of effects of plateau zokor disturbance and climate warming on species’ demography, we used mixed-effects models to relate each vital rate to treatment and/or individual size. Demographic data were pooled across plots and treatments, and then used to fit models for eight vital rates that collectively determine population dynamics: probability of survival, changes in size (i.e., growth and shrinkage), probability of reproduction, number of flowers, probability of seedling establishment, seedling size distribution, number of ramets produced per size-species plant, and ramet size distribution. Probability of survival and reproduction were modeled from the Binomial family with the logit link function. Changes in size, seedling size distribution, and ramet size distribution were modeled from the Gaussian family. Number of spikes were modeled from the negative binomial family with the log link function. Probability of seedling establishment and number of recruits produced per size-specific plant were modeled from the beta family with the logit link function (Cribari-Neto and Zeileis 2010).

To determine the best proxy for plant size in *T. lanceolata*, we used several size metrics to characterize plant survival, size changes, and reproduction. We estimated plant size using eight candidate size metrics, derived from the morphological measurements described above and either used directly or combined arithmetically (Table S2). All size metrics were also log-transformed to normalize model residuals and reduce skewness where applicable (Baudraz et al. 2025). We used the pooled demographic data pooled to construct mixed-effects models for the size-dependent vital rates, with plot included as a random effect, and assessed the overall metric of model performance using Nakagawa’s *R^2^* (Nakagawa and Schielzeth 2013, Johnson 2014, Nakagawa et al. 2017). Nakagawa’s *R^2^* includes marginal *R^2^* 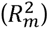 and conditional *R^2^* 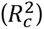 component, where 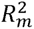 represents the variance explained by the fixed effect and 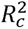 represents the variance explained by the entire model, including the random structure. Because the equations for derive *R^2^* differ depending on the model’s error distribution and the link function (Nakagawa and Schielzeth 2013), we use 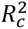 to compare metrics in their ability to explain each vital rate. The 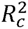 values were normalized prior to averaging them across the different vital rates (Baudraz et al. 2025):

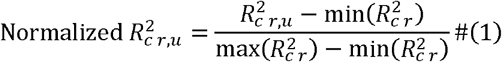

where *r* denotes the vital rate and *u* is the size metric. The variable with the highest mean normalized 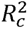 across vital rate models was selected as the proxy for plant size in each study species: log-transformed crone volume (Table S3).

To assess how plateau zokor disturbance and climate warming influence plant vital rates, we developed candidate models incorporating different combinations of fixed effects, slope variation with plant size, and random effect structures (Table S4). Models varied in whether warming and disturbance were included as main effects or interactions, and in whether the effect of plant size (slope) was constant or allowed to vary with warming, disturbance, or their interaction. Random effects were specified as random intercepts only, random slopes of plant size, or both, with or without correlations between intercepts and slopes (Ellner et al. 2016, Tredennick et al. 2018). Models were fitted using the R packages *nlme* or *glmmTMB* (Bates et al. 2015, Brooks et al. 2017). We compared candidate models using Akaike’s Information Criterion corrected for small sample size (AICc) to select the best-supported model for each vital rate (Table S5; Burnham and Anderson 2004).

### Population dynamics model

To quantify he effects of plateau zokor disturbance and climate warming on population performance, we constructed Integral Population Models (IPMs) incorporating all the demographic processes (Rees and Ellner 2009). IPMs use information on how an individual’s size influences vital rates to project population changes in discrete time (Easterling et al. 2000). Each IPM was parameterized with estimated vital rate parameters derived from the best-supported models for each vital rate (Table S6). In our IPMs, the continuous plant size was log-transformed crone volume, and the discrete time step (from *t* to *t* + 1) corresponded to one year. The size-structed IPM is

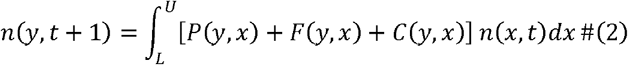

where *L* and *U* are the lower and upper bounds on the range of possible plant size for all treatments. The variable *n*(*x*, *t*) is the distribution of plant size *x* at time *t*, and *n*(*y*, *t* + 1) is the distribution of plant size *y* at time *t* + 1. The expression *P*(*y*, *x*) + *F*(*y*, *x*) + *C*(*y*, *x*) is called the kernel, *K*(*y*, *x*), which is a non-negative surface describing all possible demographic transitions from plant size *x* at time *t* to plant size *y* at time *t* + 1. The function *P*(*y*, *x*) comprises the survival-growth component of a IPM and can be decomposed into two functions that determine the probability of survival of an individual at plant size *x*, *p_s_*(*x*), and the likelihood that the individual will grow from plant size *x* to plant size *y* over a year, *g*(*y*, *x*), such that *P*(*y*, *x*) = *p_s_*(*x*) *g*(*y*, *x*). The function *F*(*y*, *x*) comprises sexual reproductive component of a IPM and can be decomposed into four functions that determine the probability of reproduction of an individual at plant size *x*, *p_f_*(*x*), the number of spikes of an individual at plant size *x*, *f_nf_* (*x*), the probability of seedling establishment *p_r_*, and the probability distribution of seedling sizes *f_ds_* (*y*), such that *F*(*y*, *x*) = *p_f_*(*x*) *f_nf_* (*x*) *p_r_ f_ds_* (*y*). The function *F*(*y*, *x*) comprises clonal reproductive component of a IPM and can be decomposed into two functions that determine the number of ramets produced per individual at plant size *x*, *f_nr_* (*x*) and the probability distribution of ramet sizes *f_dr_* (*y*), such that *F*(*y*, *x*) = *f_nr_* (*x*) *f_dr_*(*y*).

To examine the effects of plateau zokor disturbance and climate warming on long-term population viability, we calculated the population growth rate (λ) under ambient and warmed conditions across the successional stages of plateau zokor mounds. The IPM kernel was discretized into 200 × 200 matrices using the midpoint rule (Ellner and Rees 2006). Population growth rates were calculated as the dominant eigenvalues of these matrices, with λ > 1 indicating population increase and λ < 1 indicating decline toward extinction under stationary equilibrium (Caswell 1982). To quantify uncertainty around λ, we bootstrapped the demographic data 1,000 times to obtain 95% confidence intervals (CIs) under each treatment (Puth et al. 2015). Furthermore, we calculated the differences between λ values (Δλ) from each bootstrap to obtain 95% CIs of Δλ between warming, plateau zokor mounds, their interaction, and the ambient intact pasture. In addition, the aforementioned bootstrapped samples were employed in the subsequent population dynamics analysis. 写 **deltadelat**

To examine the effects of climate warming on the transient dynamics of natural populations, we quantified the demographic resilience of undisturbed populations under ambient and warmed conditions (Stott et al. 2011, Capdevila et al. 2020). We used the aforementioned matrices to estimate three components of resilience: *resistance*, *compensation*, and *recovery time* (Figure 1; Stott et al. 2012, Capdevila et al. 2022). *Resistance*, defined as the ability of a population to avoid a decrease in growth rate following disturbance, was quantified as the inverse of a population’s decrease following a disturbance relative to its undisturbed conditions. Thus, low resistance indicates large population declines relative to a stable population. *Compensation*, the ability of a population to increase its growth rate after disturbance, was quantified as the population’s increase with respect to an undisturbed population. Large increases in population size indicate high compensation, whereas small increases indicate low compensation. *Recovery time* was defined as the time required for a population to return to its stable demographic structure following disturbance. All three transient demographic resilience metrics were calculated using the *popdemo* R package (Stott et al. 2012).

To examine the relative importance of each vital rate on the population growth rate, we conducted prospective perturbation analyses, the elasticity analyses. Elasticity analysis of vital rates quantifies a proportional change in λ in response to a proportional change in vital rate (Zuidema and Franco 2001), which makes comparisons among vital rates (Griffith 2017). By conducting vital rate elasticity analyses, we were able to determine the elasticity values for survival, growth, sexual production, and clonal reproduction separately. Positive elasticity values imply that an increase in a vital rate will increase λ, which is particularly useful for distinguishing the relative importance of positive growth versus shrinkage (Zuidema and Franco 2001). To estimate uncertainty around vital rate elasticity values, we used the aforementioned bootstrapped samples to obtain their 95% CIs.

### Life table response experiment

To test how disturbance-, warming-, and their interaction-induced changes in vital rates might have led to differences in population growth rates, we conducted retrospective demographic analyses, the life table response experiment (LTRE). An LTRE analysis decomposes the observed difference in population growth rate into the contributions from different vital rates (Caswell 1989). We conducted a fixed-design LTRE on vital rate over a life cycle. The one-factor LTRE analysis follows the formulation

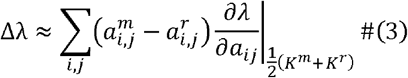

where Δλ is the difference in population growth rate between plateau zokor mounds, warming, their interaction (*m*), and the ambient intact pasture (*r*). The contribution of each vital rate is calculated as the differences between the vital rate 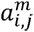 in the large transition matrix *K^m^* under treatment *m* and the corresponding vital rate 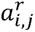 in the large transition matrix *K^r^* under reference *r*, multiplied by the sensitivity values of the mid matrix, defined as the average of *K^m^* and *K^r^* (Caswell 1989). Using the mid matrix reduces deviations arising from the nonlinear nature of vital rate sensitivities (Caswell 1982).

All analysis was conducted suing R statistical software v.4.0.5 (R Core Team 2021).

## RESULTS

### The impacts of plateau zokor disturbance and climate warming on demographic processes

To examine the effects of plateau zokor disturbance and climate warming on demographic processes of alpine herb *Thermopsis lanceolata*, we used mixed-effects models to relate vital rates to plant sizes across successional stages of plateau zokor mounds under both ambient and warmed conditions. We found that survival probability decreased with increasing plant size (χ^2^(1) = 33.39, *P* < 0.05, Table 1; Figure 4a). Warming further reduced survival across all plant sizes under ambient intact pasture (χ^2^(1) = 3.95, *P* < 0.05, Table 1; Figure 4a). In contrast, disturbance of plateau zokor increased the survival across all plant sizes under ambient condition but had only minor effects under warmed condition (Figure 4a and S2). Growth decreased with increasing plant size, whereas shrinkage increased with increasing plant size (χ^2^(1) = 47.60, *P* < 0.05, Table 1; Figure 4b, c). Warming enhanced growth but reduced shrinkage of medium-sized plants under ambient intact pasture (Figure 4b, c). Disturbance decreased growth of medium-sized plants on new mound under both ambient and warmed conditions, while increasing shrinkage of medium-sized plants on new mounds under both conditions (χ^2^(3) = 7.08, *P* = 0.07, Table 1; Figure 4b, c and S2).

**Figure 4.**
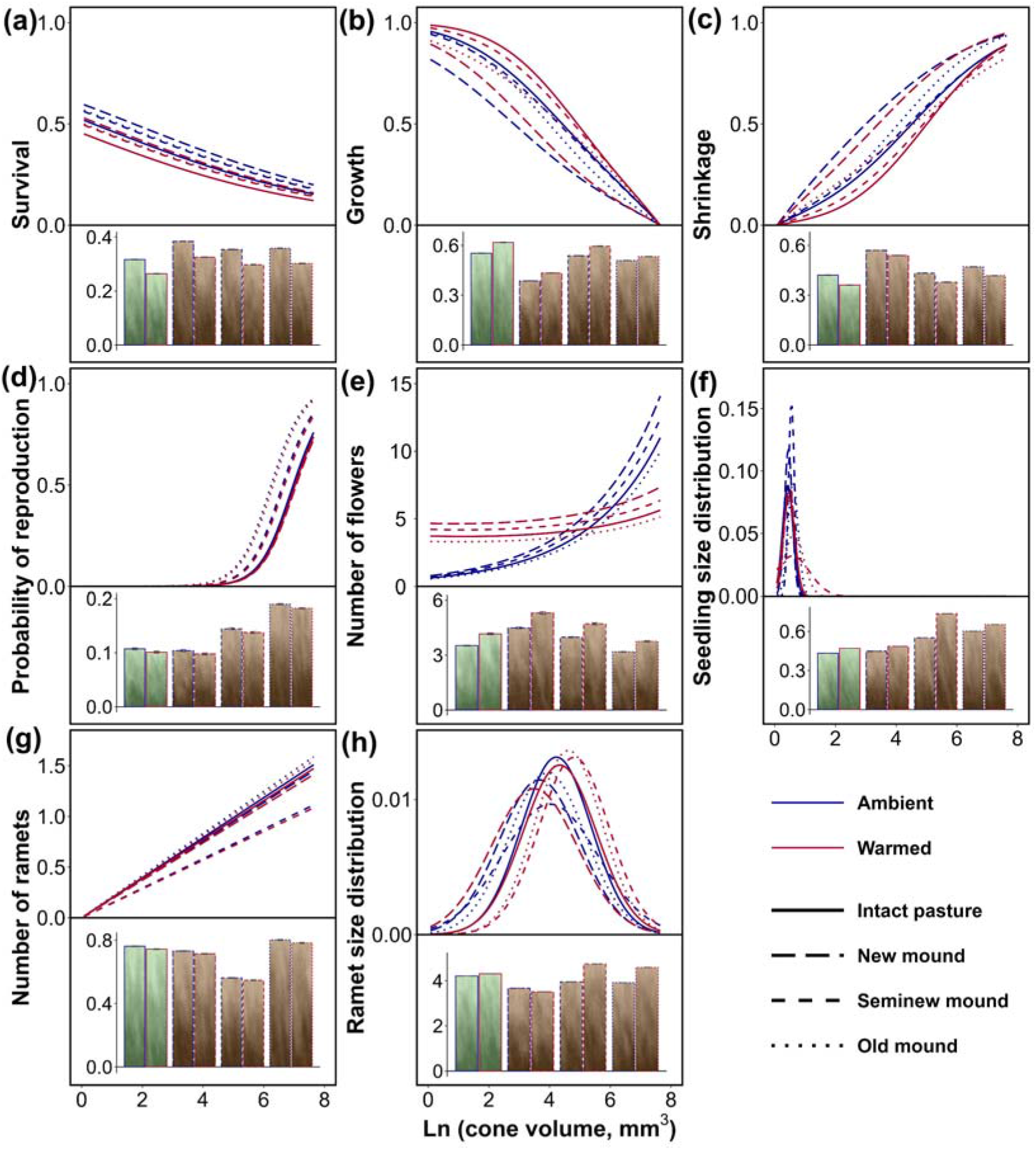
Effects of plateau zokor disturbance and climate warming on vital rates of the alpine herb *Thermopsis lanceolata* in an alpine meadow on the Tibetan Plateau. Different line types represent vital rate values across successional stages of plateau zokor mounds, whereas line colors distinguish ambient and warmed conditions. Mean values of each vital rate across all sizes are given in the bar chart at the bottom of each panel, with error bar representing 95% confidence interval.

**Table 1.** Result of analyses of fixed effects (*i.e.* warming, plateau zokor disturbance, and the natural logarithm of current plant cone volume) on each vital rate for the alpine herbs *Thermopsis lanceolata* on the Tibetan Plateau. Wald chi-squared test was provided for fixed effects in the best-supported mixed-effects models for each vital rate (Table S5), and significance was determined using type-II test.

| Parameter | $\chi^2$ | df | P | Parameter | $\chi^2$ | df | P |
| --- | --- | --- | --- | --- | --- | --- | --- |
| **Probability of survival (Binomial error, logit link)** |  |  |  | **Changes in size (Gaussian error, identity link)** |  |  |  |
| Disturbance | 3.503 | 3 | 0.320 | Disturbance | 7.075 | 3 | 0.070 |
| Warming | 3.951 | 1 | * | Warming | 1.283 | 1 | 0.257 |
| Ln (cone volume) | 33.387 | 1 | *** | Ln (cone volume) | 47.597 | 1 | *** |
| **Probability of reproduction (Binomial error, logit link)** |  |  |  | **Number of spikes (Negative binomial error, log link)** |  |  |  |
| Disturbance | 9.603 | 3 | * | Disturbance | 3.319 | 3 | 0.345 |
| Warming | 0.224 | 1 | 0.636 | Warming | 0.003 | 1 | 0.960 |
| Ln (cone volume) | 64.662 | 1 | *** | Ln (volume) | 10.438 | 1 | ** |
| - | - | - | - | Warming × Ln (cone volume) | 4.930 | 1 | * |
| **Probability of seedling establishment (Beta error, logit link)** |  |  |  | **Distribution of seedling sizes (Gaussian error, identity link)** |  |  |  |
| Disturbance | 9.057E+11 | 3 | *** | Disturbance | 9.637 | 3 | * |
| Warming | 5.432E+10 | 1 | *** | Warming | 0.539 | 1 | 0.463 |
| Disturbance × warming × | 2.866E+10 | 3 | *** | - | - | - | - |

| Number of ramets (Beta error, logit link) |  |  |  | Distribution of ramet sizes (Gaussian error, identity link) |  |  |  |
| --- | --- | --- | --- | --- | --- | --- | --- |
| Disturbance | 17.438 | 3 | *** | Disturbance | 218.475 | 6 | *** |
| Warming | 0.179 | 1 | 0.672 | Warming | 86.079 | 4 | *** |
| - | - | - | - | Disturbance × warming | 57.429 | 3 | *** |

Note: The term “Ln (cone volume)” denotes the natural logarithm of current plant volume. The term *warming* refers to ambient and warmed conditions, while the term *disturbance* refers to intact pasture, new mound, seminew mound, and old mound. The symbol × indicates an interaction between parameters. *df*, degree of freedom; \*, $P < 0.05$ ; \*\*, $P < 0.01$ ; \*\*\*, $P < 0.001$ .

The probability of reproduction and flower production increased with increasing plant size (*P* < 0.05, Table 1; Figure 4d, e). Disturbance increased reproductive probability of seminew and old mounds under both ambient and warmed conditions (χ^2^(3) = 9.60, *P* < 0.05, Table 1), while warming enhanced flower production in small plants but reduced it in large plants (Figure 4d, e and S2). Disturbance increased seedling recruitment on new mounds under both ambient and warmed conditions, but decreased seedling recruitment on seminew and old mounds under both conditions (Figure 5). Disturbance also increased the seedling size distribution of seminew and old mounds under both ambient and warmed conditions (χ^2^(3) = 9.60, *P* < 0.05, Table 1), while warming further increased seedling size distribution (Figure 4f and S2).

**Figure 5.**
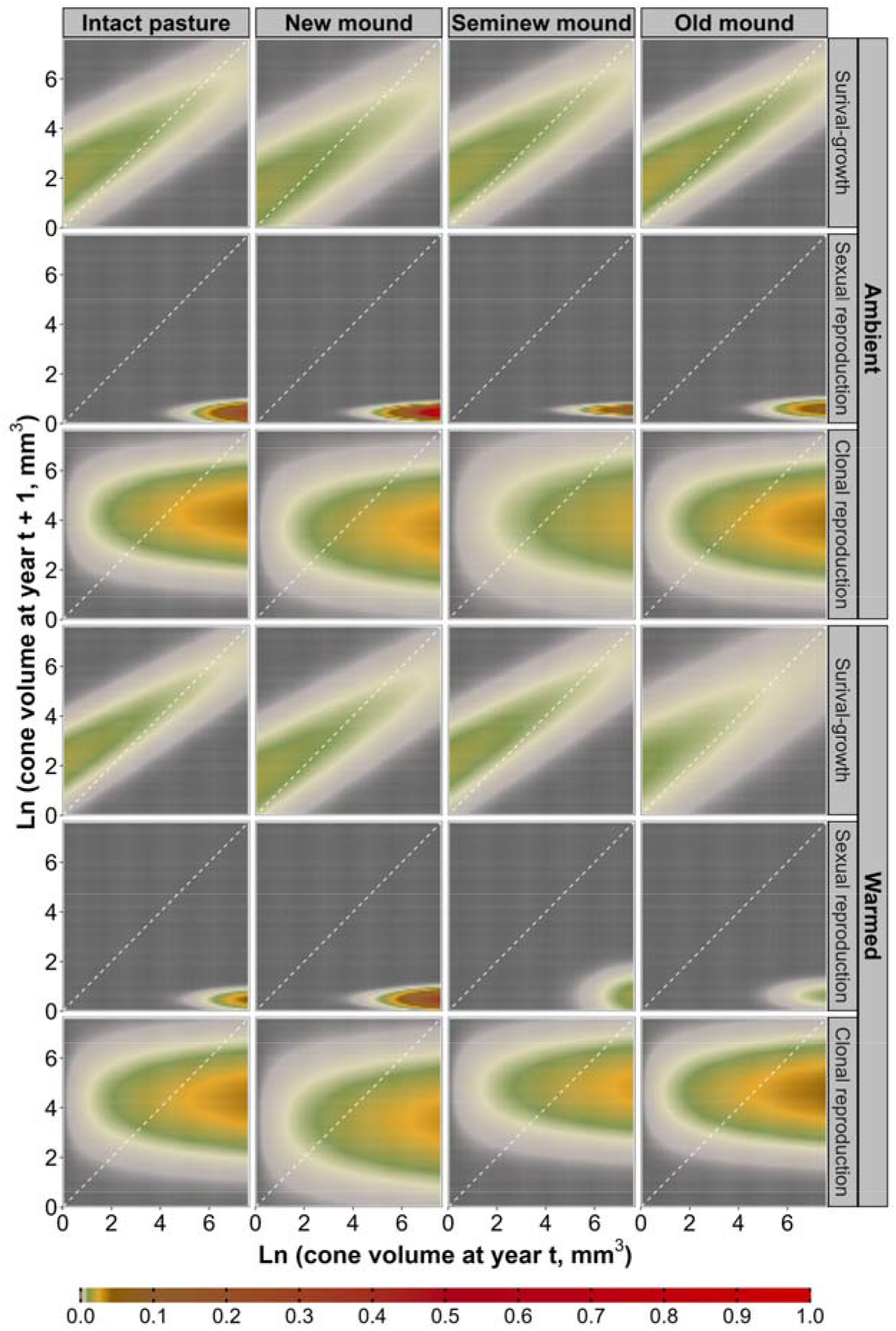
Effects of plateau zokor disturbance and climate warming on survival-growth components, sexual reproduction, and clonal reproduction of the alpine herb *Thermopsis lanceolata* in an alpine meadow on the Tibetan Plateau. The dashed 1:1 line along the diagonal represents individuals at year *t* that remained the same plant size or produced recruits of the same plant size at year *t* + 1.

Warming reduced ramet production under ambient intact pasture (Figure 4g and S2). Disturbance decreased ramet production of new and seminew mounds but increased it on old mound under both ambient and warmed conditions (χ^2^(3) = 17.44, *P* < 0.05, Table 1; Figure 4g and S2). Warming increased ramet size distribution (χ^2^(4) = 86.08, *P* < 0.05, Table 1; Figure 4h). Disturbance decreased ramet size distribution ambient condition, whereas under warmed conditions it decreased ramet size on new mounds but increased it on seminew and old mounds (χ^2^(6) = 218.48, *P* < 0.05, Table 1; Figure 4h and S2). Warming decreased the seedling recruitment under ambient intact pasture (Figure 5).

### The impacts of plateau zokor disturbance and climate warming on population growth and demographic resilience

To examine the population response to warming and plateau zokor disturbance, we built IPMs to estimate the population growth rate (λ) of *T. lanceolata* across successional stages of plateau zokor mound under both ambient and warming conditions, and quantified changes in λ relative to ambient intact pasture (Δλ). We found that the population growth rate was significantly greater than 1 under ambient intact pasture (λ = 1.138, 95% CI [1.135, 1.142]; Figure 6a). However, the population growth rate was weakened under either warmed intact pasture or disturbance from new mound, and their combined effects on population growth rate (Δλ = −0.151, 95% CI [−0.155, −0.146]; Figure 6b) was greater than the sum of their individual effects (−0.126, 95% CI [−0.131, −0.121]; Figure 6b), reducing λ below 1 (λ = 0.988, 95% CI [0.985, 0.990]; Figure 6a). Disturbance led to the population decline under seminew mound, and the superimposed effect from warming under such disturbance was greater than either individual effects but lower than the sum of their individual effects, leading to further population decline (λ = 0.925, 95% CI [0.923, 0.929]; Figure 6). Interestingly, disturbance increased the population growth rate to 1.165 under old mound, and the interactive effect of warming under such disturbance was positive and large than the disturbed effects along, further promoting the population growth (λ = 1.181, 95% CI [1.177, 1.185]; Figure 6). Additionally, warming had different effects on the transient demographic resilience metrics estimated from IPMs under intact pasture: it enhanced resistance and shortened recovery time, but weakened compensation (Table S7).

**Figure 6.**
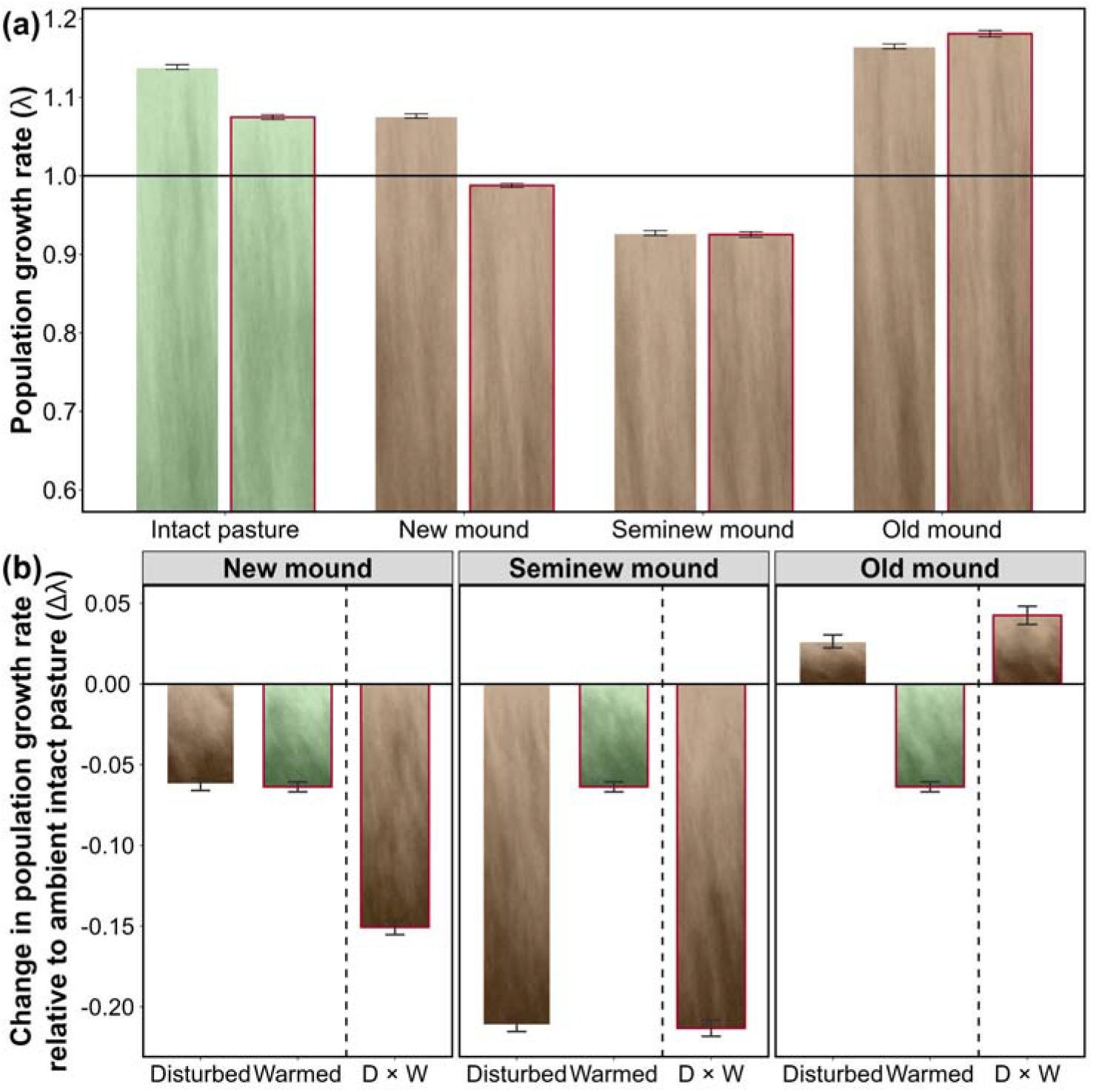
Effects of plateau zokor disturbance and climate warming on population dynamics of the alpine herb *Thermopsis lanceolata* in an alpine meadow on the Tibetan Plateau. (a) Population growth rate (λ) across successional stages of plateau zokor mound under either ambient (red lined bar chart) or warmed (unlined bar chart) conditions. (b) Changes in population growth rate between disturbed, warmed, and their interaction treatments relative to intact pasture in ambient. Error bar indicates 95% confidence interval (CI). D, disturbed treatment; W, Warmed treatment; ×, the interaction between warming and disturbance.

### The impact of plateau zokor disturbance and climate warming on vital rate elasticity

To examine how the importance of vital rate for population maintenance response to plateau zokor disturbance and climate warming, we conduced elasticity analyses of vital rate on the population growth rate. We found that the population growth rate was overwhelming most elastic to changes in clonal reproduction and survival across successional stages of plateau zokor mound under both ambient and warmed conditions (Figure 7). The elasticities of clonal reproduction increased under warmed intact pasture but decreased across successional stages of mounds under ambient conditions, whereas the survival elasticities showed the opposite pattern (Figure 7). Importantly, the combined effects of warming and disturbance on clonal reproduction elasticity were smaller than the additive effects of warming and new mound, larger than their addition effects from warming and seminew or old mound (Figure S3). Additionally, the elasticities to changes in plant size (i.e. positive growth) and sexual reproduction were positive but much smaller under all treatments (Figure 7). However, the elasticity to shrinkage was negative in all populations, indicating a negative effect on population maintenance as a result of individual become smaller (Figure 7).

**Figure 7.**
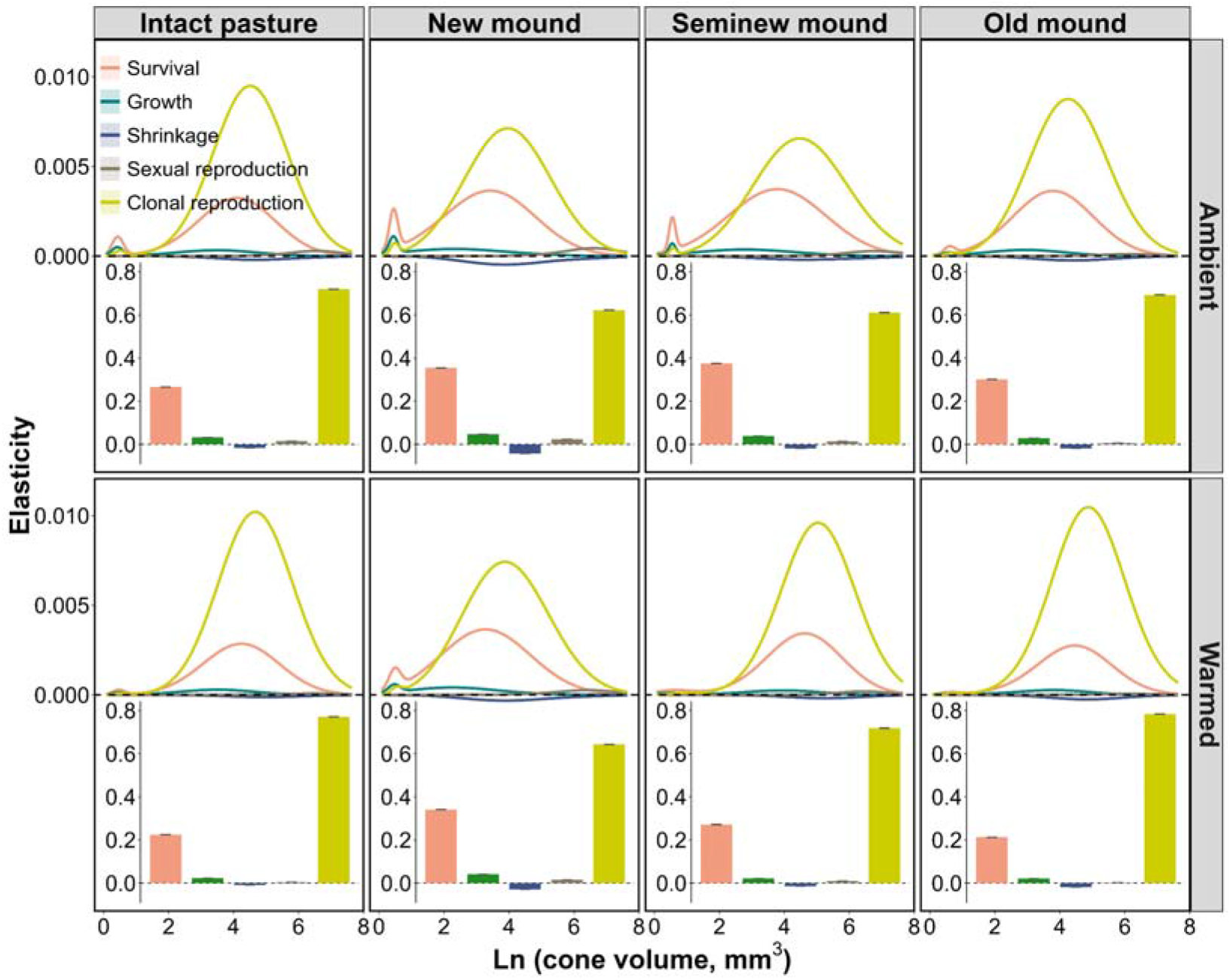
Effects of plateau zokor disturbance and climate warming on vital rate elasticity of the alpine herb *Thermopsis lanceolata* in an alpine meadow on the Tibetan Plateau. Cumulative elasticities of each vital rate across all sizes are given in the bar chart at the bottom of each panel, with error bars representing 95% confidence intervals.

### The impact of plateau zokor disturbance and climate warming on vital rate contribution

To examine how disturbance-, warming-, and their interaction-induced changes in λ were caused by changes in vital rates, we conducted the LTRE analysis. We found that warming-induced small reduction in λ of intact pasture was mainly caused by decreases in survival and ramet production of medium-sized plants (Figure 8). In contrast, the reduction in λ of new mound under both ambient and warmed conditions was primarily driven by decreases in large ramet size distribution, medium ramet production, and changes in medium plant sizes (i.e. growth and shrinkage) (Figure 8). The greater reduction in λ under warmed new mound compared to the additive expectation was mainly due to a larger decrease of ramet size distribution (54%; Figure 8, S4, and S5). The population decline on ambient seminew mound was primarily attributed to a large decrease of medium ramet production and a small decrease of medium ramet size distribution, whereas the population decline on warmed seminew mound was mainly caused by a large decrease of medium ramet production and a small decreased survival of medium plants (Figure 8). The smaller reduction in λ under warmed seminew mound relative to additive effects was mainly caused by a larger increase in ramet size distribution (70%; Figure 8, S4 and S5). The small increase in λ on ambient old mound was largely due to increases in ramet production and survival of medium plants, while the large increase in λ under warmed old mound was mainly caused by increases in large ramet size distribution and medium ramet production (Figure 8). The larger increase in λ under warmed old mound relative to additive effects was mainly caused by larger increased ramet size distribution (73%; Figure 8, S4 and S5).

**Figure 8.**
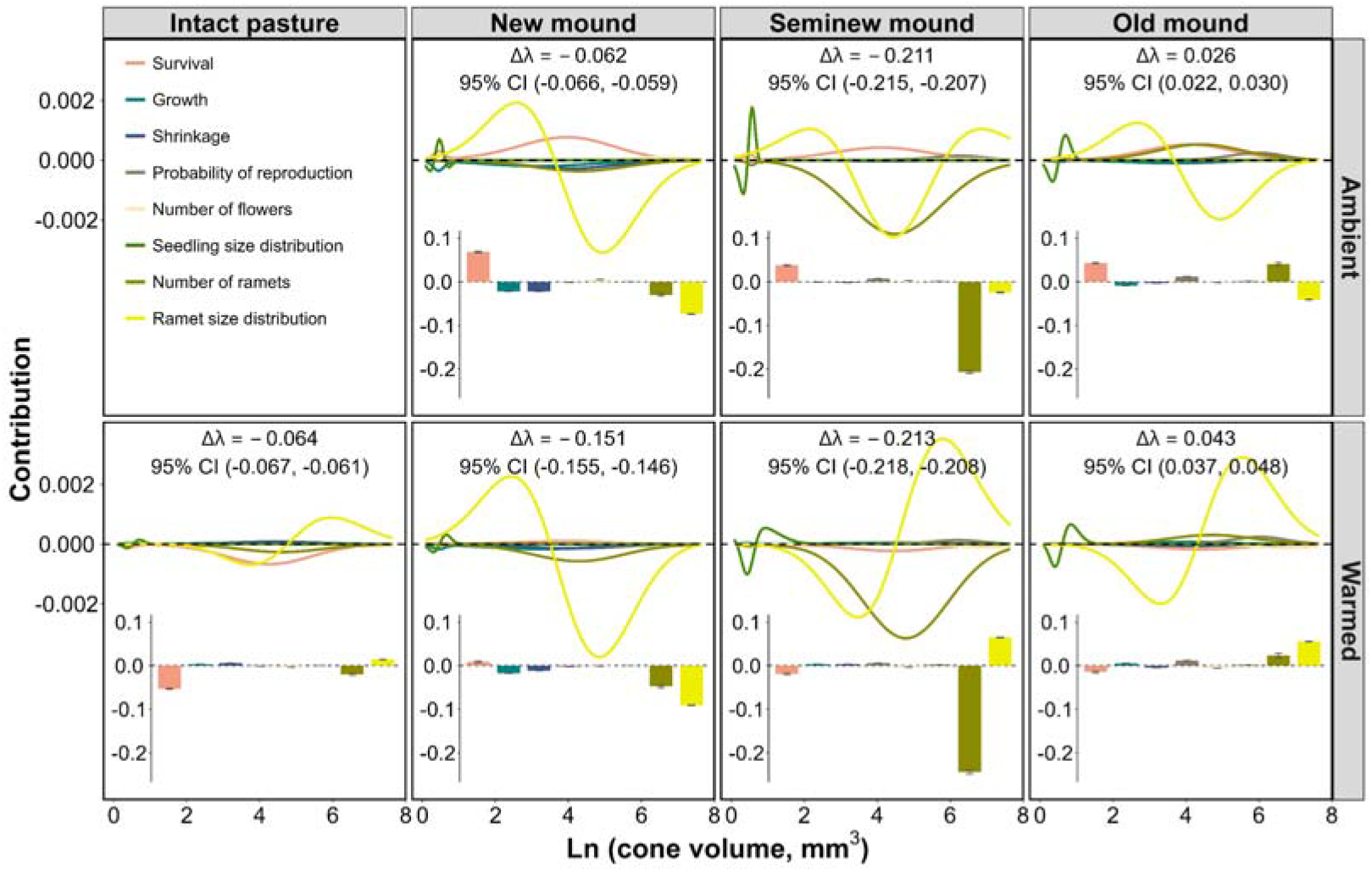
Results of life table response experiment analysis to examine the effects of plateau zokor disturbance and climate warming on vital rate elasticity of the alpine herb *Thermopsis lanceolata* in an alpine meadow on the Tibetan Plateau. Values showed vital rate contributions to differences in population growth rates between warmed, disturbed, and their interaction treatments relative to intact pasture (Δλ). Cumulative contributions of each vital rate across all sizes are given in the bar chart at the bottom of each panel, with error bars representing 95% confidence intervals.

## DISCUSSION

Quantitatively assessing plant population dynamics following disturbance under current and warming climates is crucial for biodiversity conservation and management decision-making (Huntly and reichman 1994, Liu et al. 2018). Such assessments require long-term demographic studies that capture the population performance across post-disturbance recovery processes (Easterling et al. 2000, Caswell 2001). However, few studies have explicitly quantified post-disturbance demographic responses, particularly under climate warming. Here, using a space-for-time substitution design across different recovery stages of mounds created by plateau zokors (Pickett 1989, Blois et al. 2013, Niu et al. 2020), we examined the demographic responses of a dominant alpine herb (*T. lanceolatum*) to plateau zokor disturbance under ambient and *in situ* active warming conditions on the Tibetan Plateau. We found that the demographic effects of plateau zokor disturbance on *T. lanceolatum* were strongly stage-dependent under both ambient and warmed conditions: population growth rates initially decreased on new mounds, further declined on seminew mounds, but eventually exceeded initial levels on old mounds, suggesting a shift from short-term demographic limitation to long-term demographic enhancement during post-disturbance succession. This stage-dependent response was largely driven by clonal reproduction (i.e., clonal production and/or ramet size distribution), which emerged as both the most sensitive vital rate and the primary contributor to variation in population growth rates across recovery stages. These findings align with theoretical predictions and previous empirical studies showing that high variation in key vital rates can constrain population growth under disturbance yet facilitate recovery once conditions stabilize (McDonald and Jones 2002, Koons et al. 2009, Jongejans et al. 2010). Together, these findings indicate that stage-dependent demographic responses allow disturbed populations to compensate for early losses and recover over the course of succession, with clonal reproduction playing a central role in mediating these effects under both current and warming climates. These stage-dependent demographic responses align with classical disturbance–succession theory, illustrating how demogrpahic processes, such as clonal reproduction, can shape successional trajectories at the community and ecosystem levels (Pickett and White 1985, Standish et al. 2014, Ingrisch and Bahn 2018).

The combined effects of disturbance and warming on demography may differ from the sum of their individual effects (Cote et al. 2016, Zettlemoyer 2023). We found that the combined effects of plateau zokor disturbance and warming on population growth rates of new mounds was greater than the sum of their individual effects, leading to population decline on new mounds. Such synergistical effects was mainly due to a larger decrease of ramet size distribution. Secondly, we found that the combined effects of plateau zokor disturbance and warming on population growth rates of seminew mounds was greater than either individual effects but lower than the sum of their individual effects, leading to further population decline. The smaller reduction in λ under warmed seminew mound relative to additive effects was mainly caused by a larger increase in ramet size distribution. Lastly, combined effects of plateau zokor disturbance and warming was positive and large than the disturbed effects along, further promoting the population growth. The larger increase in λ under warmed old mound relative to additive effects was mainly caused by larger increased ramet size distribution. ramet size distribution. Climate warming may enhance the demographic resilience.

---

**Table S1.** Demogrpahic resilience (compenstion, resistance, and recovery time) of the alpine herb *Thermopsis lanceolata* under ambient and warmed conditions in an alpine meadow on the Tibetan Plateau.

| Demographic resilience | Ambient | Warmed | $\Delta_{\text{warmed} - \text{Ambient}}$ |
| --- | --- | --- | --- |
| Compensation | 3.068 [3.037, 3.099] | 1.736 [1.728, 1.744] | -1.332 [-1.362, -1.302] |
| Resistance | 0.311 [0.309, 0.313] | 0.295 [0.293, 0.297] | -0.016 [-0.017, -0.015] |
| Recovery time | 1.412 [1.407, 1.418] | 1.177 [1.173, 1.182] | -0.235 [-0.239, -0.231] |

Note: values in square brackets are 95% confidence intervals of demographic resilience.

